# VLP-Based Model for Study of Airborne Viral Pathogens

**DOI:** 10.1101/2024.01.03.574055

**Authors:** Michael Caffrey, Nitin Jayakumar, Veronique Caffrey, Varada Anirudan, Lijun Rong, Igor Paprotny

## Abstract

The recent COVID-19 pandemic has underscored the danger of airborne viral pathogens. The lack of model systems to study airborne pathogens limits the understanding of airborne pathogen distribution, as well as potential surveillance and mitigation strategies. In this work, we develop a novel model system to study airborne pathogens using virus like particles (VLP). Specifically, we demonstrate the ability to aerosolize VLP and detect and quantify aerosolized VLP RNA by Reverse Transcription-Loop-Mediated Isothermal Amplification (RT-LAMP) in real-time fluorescent and colorimetric assays. Importantly, the VLP model presents many advantages for the study of airborne viral pathogens: (i) similarity in size and surface components; (ii) ease of generation and noninfectious nature enabling study of BSL3 and BSL4 viruses; (iii) facile characterization of aerosolization parameters; (iv) ability to adapt the system to other viral envelope proteins including those of newly discovered pathogens and mutant variants; (v) the ability to introduce viral sequences to develop nucleic acid amplification assays.

**Importance:** Study and detection of airborne pathogens is hampered by the lack of appropriate model systems. In this work we demonstrate that noninfectious Virus Like Particles (VLP) represent attractive models to study airborne viral pathogens. Specifically, VLP are readily prepared, are similar in size and composition to infectious viruses, and are amenable to highly sensitive nucleic acid amplification techniques.

## Introduction

### Outbreak of COVID-19

In December of 2019, a novel coronavirus (CoV) outbreak in the Wuhan region of China became evident, which was subsequently called SARS-CoV-2 virus, the causative agent of COVID-19 [1,2]. To date, >500 million infections with >6 million deaths have been documented. Notably, the number of COVID-19 cases quickly surpassed that of the 2003-2004 SARS-CoV outbreak [3] and the virus has become the most-deadly pandemic in >100 years. Beyond the obvious costs to human health and life, COVID-19, for which humans have little or no pre-existing immunity, has very significant social and economic costs. Indeed, during the first two years of the pandemic significant regions of the world were in full or partial lockdown situations with the goal of mitigating the spread of the virus and not overwhelming the national health systems [4]. Finally, it is evident that SARS-CoV-2 is primarily transmitted via aerosols [5,6], which can be suspended for hours, and thus SARS-CoV-2 is considered an airborne viral pathogen.

### Other airborne viral pathogens

The COVID-19 pandemic underscores the danger that airborne viral pathogens pose to humanity. Notably, the pandemic H1N1 influenza outbreak of 1918 resulted in over 50 million deaths worldwide with a fatality rate of ∼3% [7,8]. Despite improved vaccination efforts and better treatments, seasonal influenza is still responsible for greater than 250,000 deaths per year worldwide [7,9]. Moreover, the recent outbreaks of avian strains harboring HA types H5 or H7, which exhibit mortality rates of >40% (i.e. >10X that of the 1918 strain) [10] are particularly concerning. Fortunately, avian influenza strains have not readily adapted for human-to-human transmission and thus to-date human fatalities have been relatively rare; however, adaptation of avian strains for human transmission occurs naturally [11]. Taken together, coronaviruses and influenza viruses, as well as other respiratory viruses, pose pandemic risks with potentially high mortality rates [5]. Consequently, airborne viruses clearly warrant continued efforts for surveillance, mitigation strategies, and the development of efficacious therapeutics and vaccines.

### Limitations of current models to study airborne viral pathogens

Bioaerosols have been intensely studied in the laboratory [12,13]. Due to biosafety concerns, many airborne viral pathogens present challenges for laboratory-based studies. For example, SARS-CoV-2 and avian influenza strains are studied under Biosafety Level 3 (BSL3) conditions. An alternative approach for study of viral pathogens under BSL1 and BSL2 conditions, which are standard to most laboratories, would be viruses inactivated by heat, ultraviolet light, or gamma irradiation (e.g. the studies of aerosolized SARS-CoV-2 in [14]). However, BSL3 facilities are often required to generate the inactivated viruses and thus samples are limited in quantity and availability. In addition, treatment by heat denatures proteins present and thus the surface properties of the virus may be negatively affected and detection by antibodies may be compromised. Alternatively, ultraviolet light and gamma irradiation treatments result in damage to the viral RNA/DNA, which may preclude detection by nucleic acid amplification techniques or subsequent sequencing efforts. On the other hand, bacteriophages such as MS2 and PR772 are popular models of airborne pathogens due to their biosafe nature, facile preparation, ease of aerosolization due to their relatively small size (20-200 nm in diameter), and detection by standard nucleic acid amplification techniques or cell culture [15]. However, there are a number of limitations with the use of aerosolized bacteriophages. First, the size of the phage is often significantly smaller than the virus of interest (e.g. MS2 phage diameter is ∼25 nm and SARS-CoV-2 diameter is ∼120 nm). Furthermore, many of the popular phages have a very different surface composition, which may be expected to affect aerosolization. For example, phages MS2 and PR772 are unenveloped and thus lack the lipid membrane present in enveloped viruses such as influenza and coronavirus. Accordingly, we propose to develop an alternative model system with attractive biosafety and biological profiles that could be used in studies of airborne viral pathogens. In what follows we describe our efforts to exploit VLP as models for the study of airborne viral pathogens, with a first focus on SARS-CoV-2.

## Materials and Methods

### VLP Generation and Characterization

Plasmids containing SARS-CoV-2 spike and the packaging construct pNL4-3.Luc.R-E− were co-transfected into 293T cells, as previously described by our laboratories [16–18]. VLP were quantified by a LFA assay for HIV p24 (GoStix Plus, Takara). In this case the number of virions per volume is calculated by the approximate stoichiometry of p24 in VLP (∼2000/virion). The presence of SARS-CoV-2 spike was verified by a Western Blot analysis using the monoclonal antibody CR-3022 (BEI Resources) and comparison to heat inactivated and gamma irradiated SARS-CoV-2 (BEI Resources), with equivalent amounts of total protein used.

### LAMP and RT-LAMP

The positive control sample consisted of the pNL4-3.Luc.R-E-plasmid, which contains the HIV pol gene, and is used as a packaging vector for the formation of VLP [19]. Plasmid concentration was determined by the absorbance at 260 nm using a ThermoScientific NanoDrop One spectrophotometer and a plasmid molecular weight of 9.2×10^6^ Da. Unless noted, VLP samples were treated by incubation at 95 °C for 10 minutes prior to analysis. The LAMP and RT-LAMP assays were performed using the NEB WarmStart^®^ Colorimetric LAMP 2X Master Mix (DNA & RNA) (#M1800) at 65 °C for 40 min [20]. Real-time LAMP and RT-LAMP were performed using the Bio-Rad CFX Duet PCR System in the 96 well format with 10 or 20 ul samples. Fluorescence changes were detected by the addition of SYBR Green to the reaction mixtures before running the assay (Invitrogen). Primers are generated by IDT based on the work of Curtis et al. [21]: F3: 5’-AGTTCCCTTAGATAAAGACTT-3’, B3: 5’-CCTACATACAAATCATCCATGT-3’, FL: 5’-GGTGTCTCATTGTTTATACTA-3’, BL: 5’-GCATGACAAAAATCTTAGA-3’, FIP: 5’-GTGGAAGCACATTGTACTGATATCTTTTTGGAAGTATACTGCATTTACCAT-3’, BIP: 5’-GGAAAGGATCACCAGCAATATTCCTCTGGATTTTGTTTTCTAAAAGGC-3’.

### Aerosolization experiments

The aerosolization experiments were performed with a custom built cabinent in a biosafety cabinet located in a BSL2 laboratory. Aerosolization of VLP utilized the Bio-Aerosol Nebulizing Generator (BANG) from CH Technologies in the Multi Pass Atomization (MPA) mode. The pressurized air flow into the nebulizer was 4.72 l/min and the vacuum pump flow rate for collection on the filter was 225 ml/min. Based on p24 concentration, the VLP solution concentration was estimated to be 3.2 × 10^9^ virions/ml in PBS buffer. Aerosolized droplets were collected using Millipore 13 mm 0.45 µm MCE filters. Bioaerosol concentration, which is a combination of water droplets and VLPs, was estimated to be 1.33 mg/m^3^ using the TSI DustTrak II optical particle counter. Quantification of VLPs deposited on the filter were estimated from successive dilutions in the RT-LAMP reaction using an LOD of 5-30 copies of viral RNA, based on the plasmid and VLP experiments containing known amounts of DNA or RNA.

## Results and Discussion

### Rationale for the VLP model

Virus Like Particles (VLP), which lack the genetic material for productive infection and are typically derived from co-transformation of plasmids containing packaging, reporter genes and/or surface proteins, have been used as surrogates for studying infectious viruses for many years [22]. VLP present numerous advantages with respect to infectious viruses including: (i) relatively safe biosafety profiles; (ii) ease of generation; (iii) similarity in size and surface composition; (iv) the ability to protect and deliver RNA. Previous uses of VLPs include detailed studies of viral entry mechanisms [16], the use of VLP to discover and characterize novel therapeutics [17,18], use as vaccine vectors, and drug or nucleic acid delivery systems [15,23].

### Generation of VLP Bearing SARS-CoV-2 Spike

As a first step, we generated VLP using the HIV vector pNL43R-E-as the packaging vector and an expression plasmid containing SARS-CoV-2 spike. The packaging vector contains the entire HIV genome with 2 mutations that result in nonfunctional envelope and rec proteins and thus the VLP are considered noninfectious and produced under BSL2 conditions that are standard to many laboratories. One advantage of this particular packaging vector is that the resulting VLP contain a copy of the viral RNA of the HIV genome, which we will exploit as a marker to track aerosolized virus by nucleic acid amplification. Addition of the second plasmid containing SARS-CoV-2 spike in the transfection results in VLP containing an authentic version of the viral envelope protein in the VLP membrane. Consequently, the VLP spike are almost identical in size (∼100 nm, [24]) and surface composition (spike and membrane lipids) to infectious SARS-CoV-2. As shown by **Fig. 1**, Western blot analysis of VLP, heat inactivated SARS-CoV-2, and gamma irradiated SARS-CoV-2 show the presence of spike in all 3 samples (we attribute the lower levels of spike observed in the heat inactivated sample to be due to the use of a conformation dependent monoclonal antibody).

**Figure 1:**
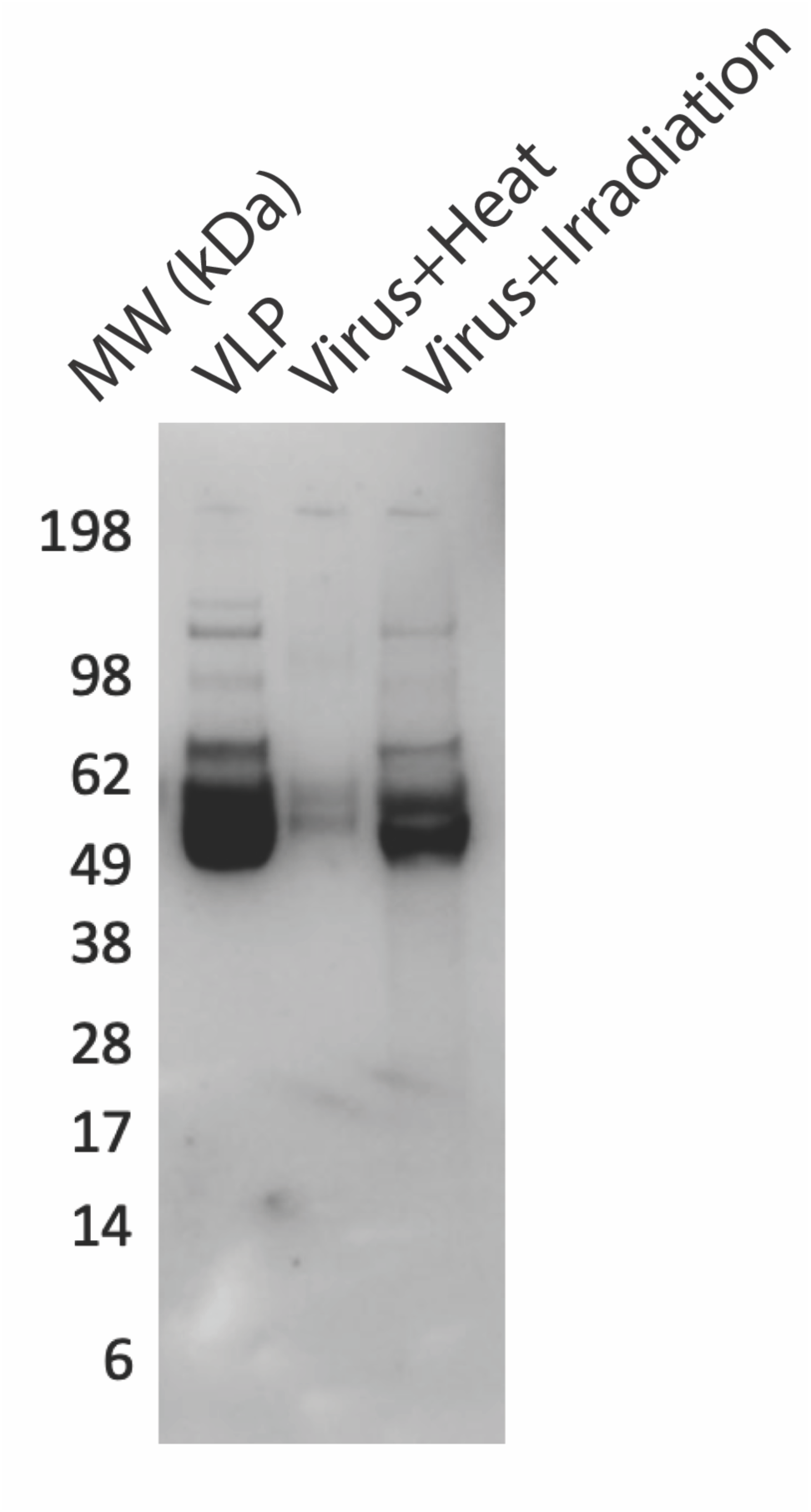
Western blot analysis for presence of SARS-CoV-2 spike in VLP and inactivated SARS-CoV-2. For this analysis, equivalent amounts of total protein were used.

### Development of the LAMP Assay Using Plasmid DNA

In the second step, we developed a nucleic acid amplification assay to track aerosolized VLP. Previously, Curtis et al. [21] developed a LAMP-based diagnostic test for HIV using a set of 6 primers to the HIV pol gene. We first tested the ability of the synthesized primer set to detect the HIV pol gene contained in the packaging vector (c.f. Materials and Methods) using the pNL43R-E-plasmid. As shown in **Fig. 2a** the Real Time LAMP assay shows fluorescence increases in a time and concentration dependent manner. Moreover, the concentration (copy number) dependence of the threshold time (Cq) exhibits the expected logrithmic relationship (**Fig. 2b**), supporting the ability to quantify the amount of nucleic acid present. Interestingly, the slope of the relationship suggests an efficiency of 280% (Efficiency=10^(−1/slope)^-1). Notably, efficiencies higher than 100% are often observed in LAMP assays [25–27]. In the next step we tested the ability to use a the colorimetric LAMP assay, which is based on the sensitivity of the dye phenol red to pH changes resulting from nucleic acid amplification [20]. As shown in **Fig. 3a**, the colorimetric assay shows the expected transition from pink to yellow over the temperature range of 62-68 °C. Moreover, using successive dilutions of the template (pNL43R-E-) we estimate that the limit of detection (LOD) is <12.5 gene copies for the colorimetric assay (**Fig 3b**) and <6.75 gene copies for the fluorescence assay (**Fig 3c**).

**Figure 2:**
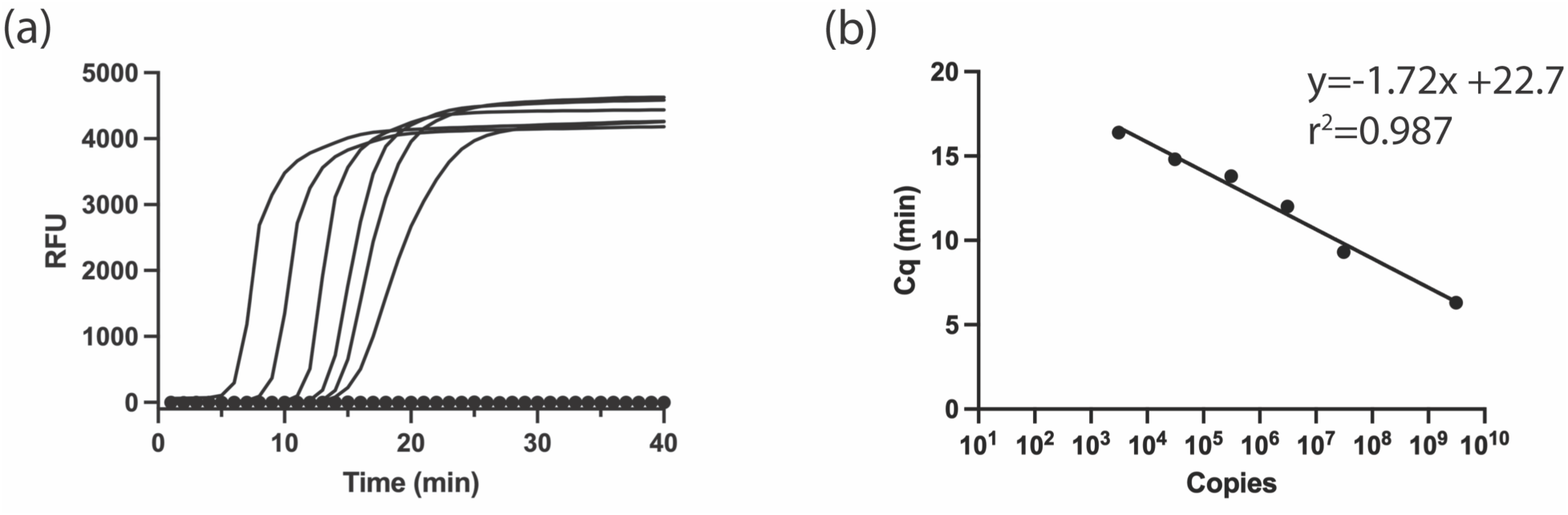
Real-time quantitative fluorescent LAMP analysis to detect plasmid. (a) Amplification plots of solutions containing 3.1 × 10^9^, 3.1 × 10^7^, 3.1 × 10^6^, 3.1 × 10^5^, 3.1 × 10^4^, and 3.1 × 10^3^ gene copies (left to right). The control experiment, which contains no template, is shown as closed circles. (b) Standard curves for the Cq values versus gene copy number. Plasmid pNL4-3.Luc.R-E-. was used for these experiments.

**Figure 3:**
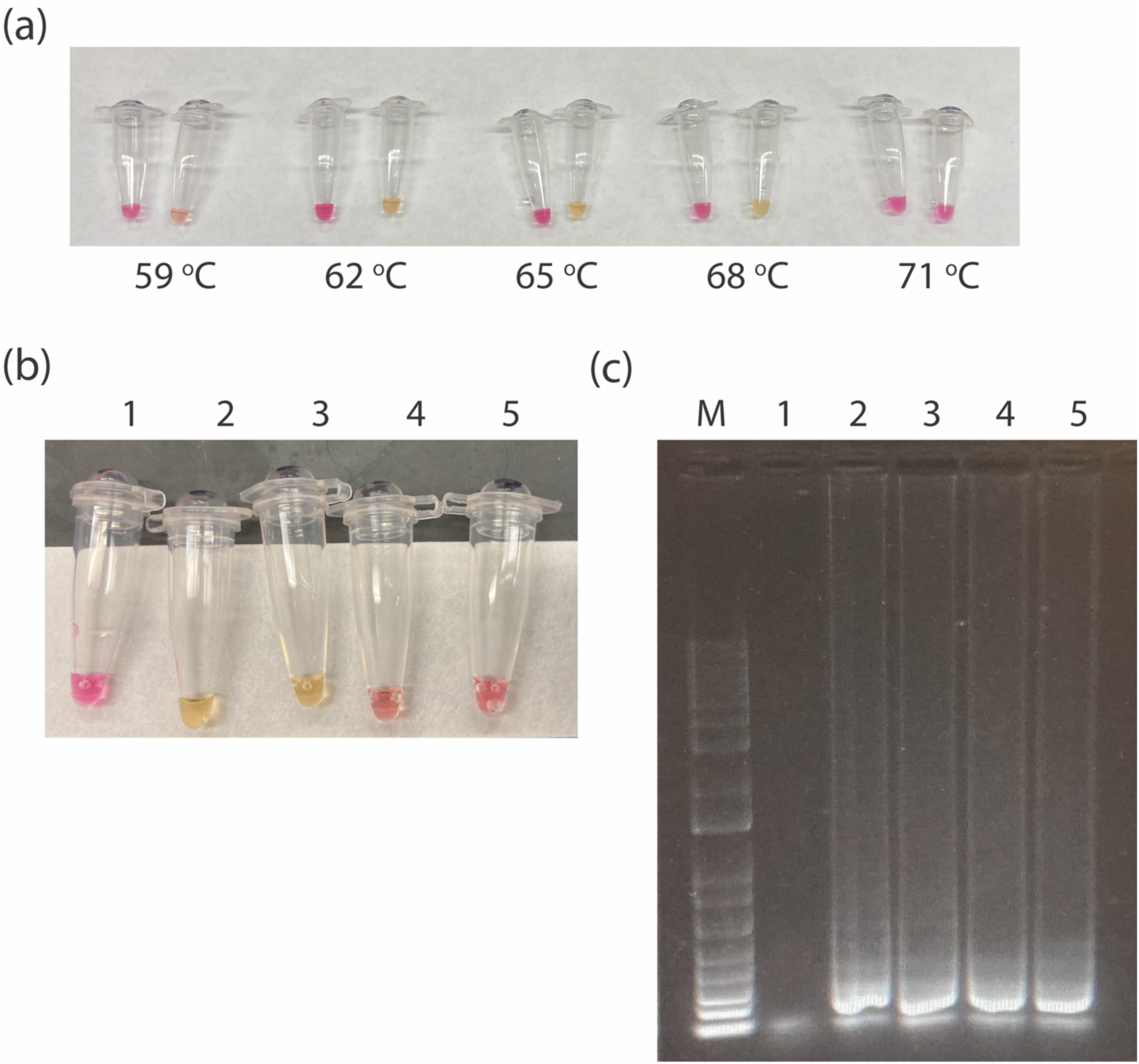
Temperature and sensitivity limits for detection by LAMP. (a) LAMP colorimetric assay for different temperatures (no template control on the left and added plasmid on the right). (b) Colorimetric LAMP assay. (c) SYBR Green stained agarose gel of LAMP reaction products. M=Marker, 1=NTC, 2=50 copies, 3=25 copies, 4=12.5 copies, 5= 6.75 copies.

### Detection of Viral RNA in VLP

As noted above, the VLP used in this study contain ∼1 copy of the HIV genome in the form of RNA, which includes the HIV pol gene. Consequently, in the next step we performed a RT-LAMP assay to detect viral RNA present in VLP. First we tested the necessity for pre-treatment of the VLP to liberate the viral RNA. As shown by the real-time RT-LAMP assay in **Fig. 4**, pre-treatment of the VLP by heat or detergent+heat do not significantly change the sensitivity (i.e. amount of viral RNA detected). Specifically, the Cq for untreated, heat treated, or detergent+heat treated VLP was 11.5 ± 0.4, 10.4 ± 0.2 and 10.4 ± 0.2 minutes, respectively. In the next step, we assessed the sensitivity of the assay using a series of VLP dilutions of stock concentrations quantified by the ELISA assay for p24. As shown by the colorimetric RT-LAMP assay and the SYBR green stained agarose gel of the RT-LAMP reaction, we estimate the LOD to be <30 gene copies of RNA (**Fig 5ab**).

**Figure 4:**
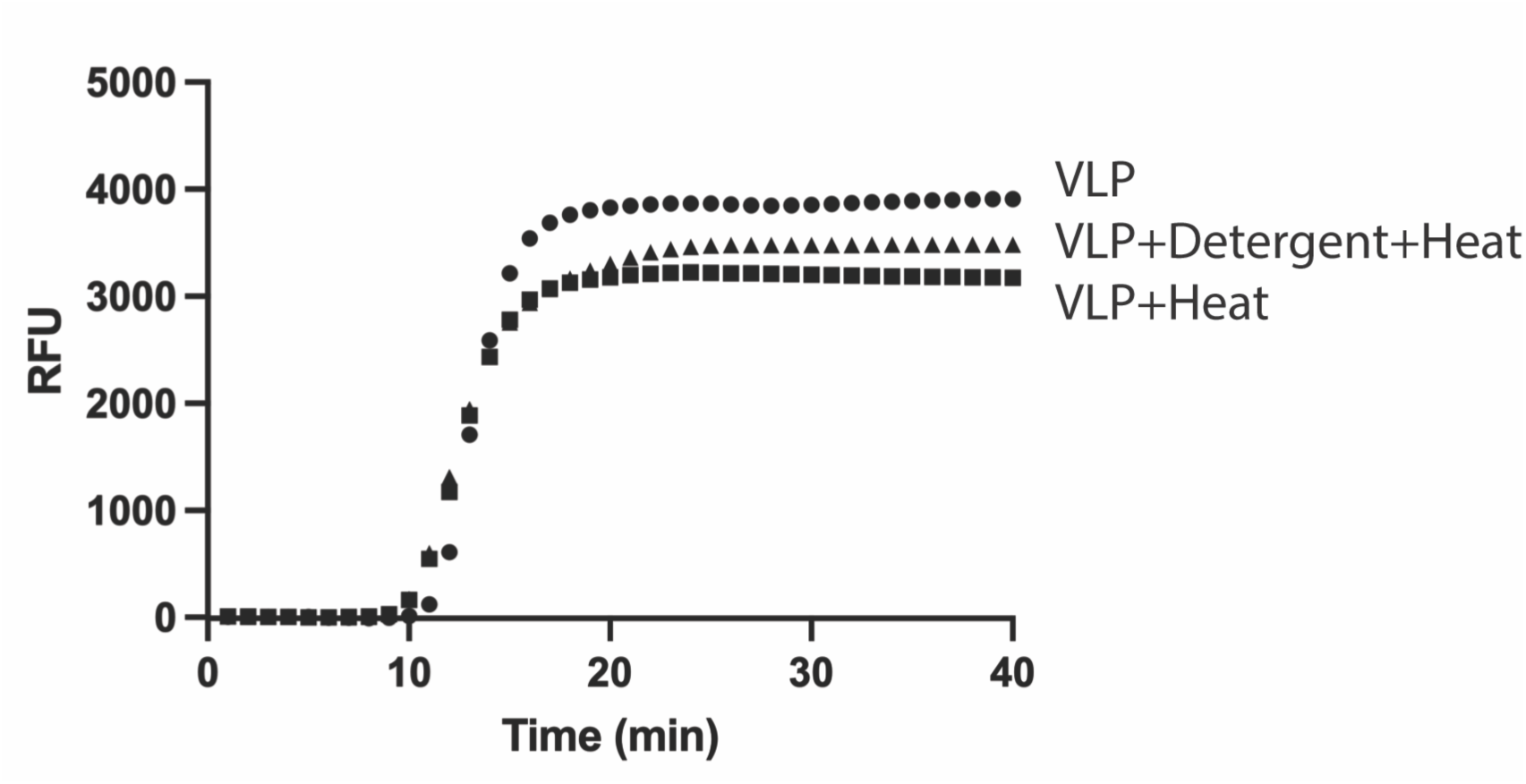
Real-time quantitative fluorescent Reverse-Transciption LAMP. For these experiments, heat treatment corresponded to incubation of VLP at 95 °C for 10 minutes before analysis and detergent treatment corresponded to VLP dilution into a buffer containing 12.5 mM TCEP, 5 mM EDTA, and 0.002% Triton X100 at pH 8.0.

**Figure 5:**
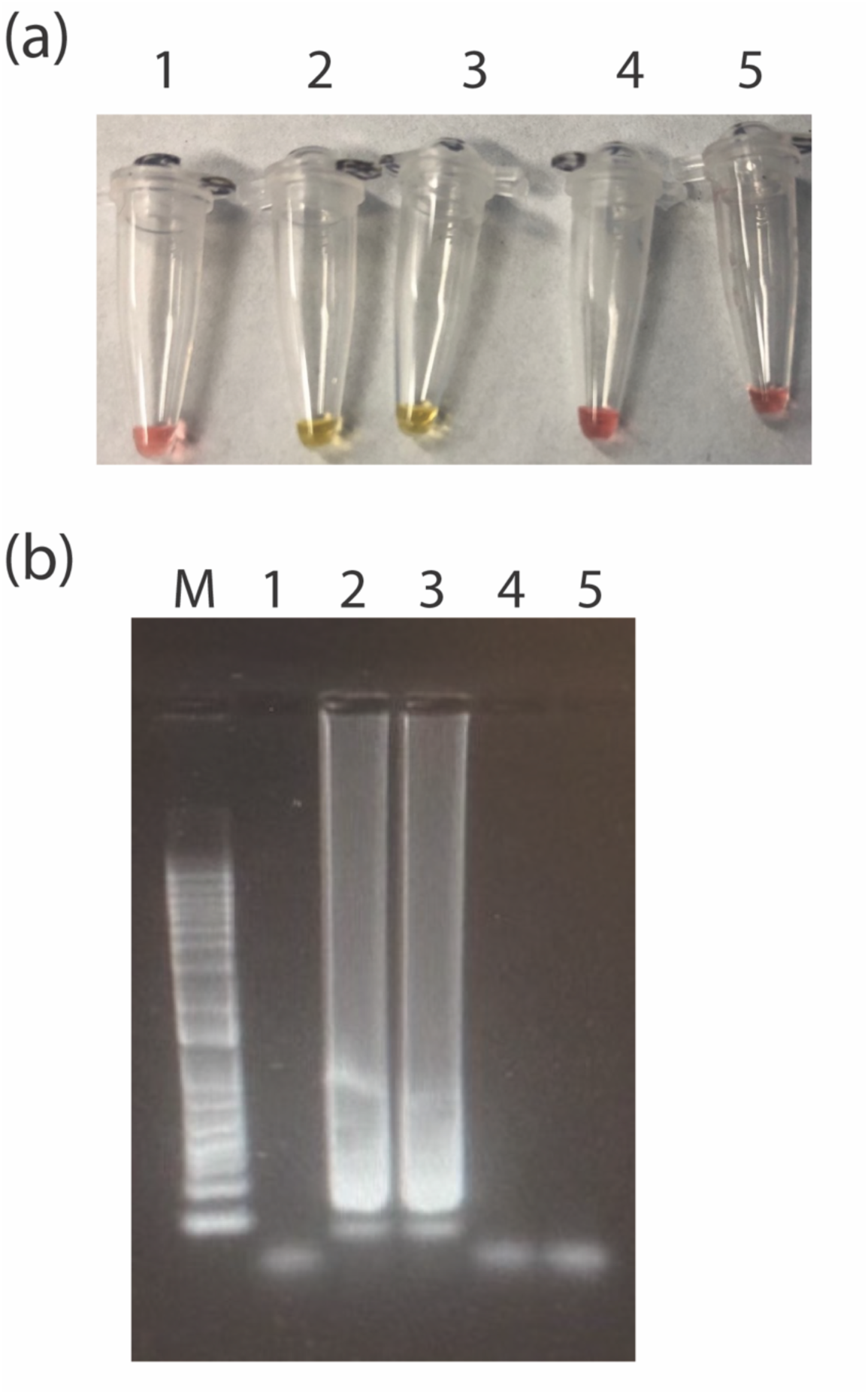
Detection of VLP by RT-LAMP. (a) Colorimetric RT-LAMP assay. (b) SYBR Green stained agarose gel of LAMP reaction products. M=Marker, 1=NTC, 2=300 copies, 3=30 copies, 4=3 copies, 5= 0.3 copies.

### Aerosolization and Detection of VLP Viral RNA

In the final step, we designed a biosafety cabinet to study aerosolized VLP. As shown in **Fig 6**, the principal components of the cabinet consist of a pressurized air inlet, a Collison nebulizer, and a filter to collect aerosolized VLP. Placing the biosafety cabinet in a biosafety hood located in our BSL2 laboratory, we then aerosolized VLP and collected the aerosolized VLP on a MCE filter (0.45 μM). Subsequently, an extract of the filter was prepared and analyzed for the presence of VLP RNA by RT-LAMP using the assay described above. As shown in **Fig 7a**, the colorimetric RT-LAMP assay suggests the presence of HIV pol gene contained in the VLP RNA in samples 1 and 2, which correspond to undiluted extract and a 1:10 extract dilution. Further analysis by a SYBR Green stained agarose gel confirms the presence of amplified nucleic acid in samples 1 and 2 (**Fig 7b**). Consequently, we have shown the ability to collect aerosolized VLP containing SARS-CoV-2 and detect the RNA contained in the VLP by RT-LAMP.

**Figure 6:**
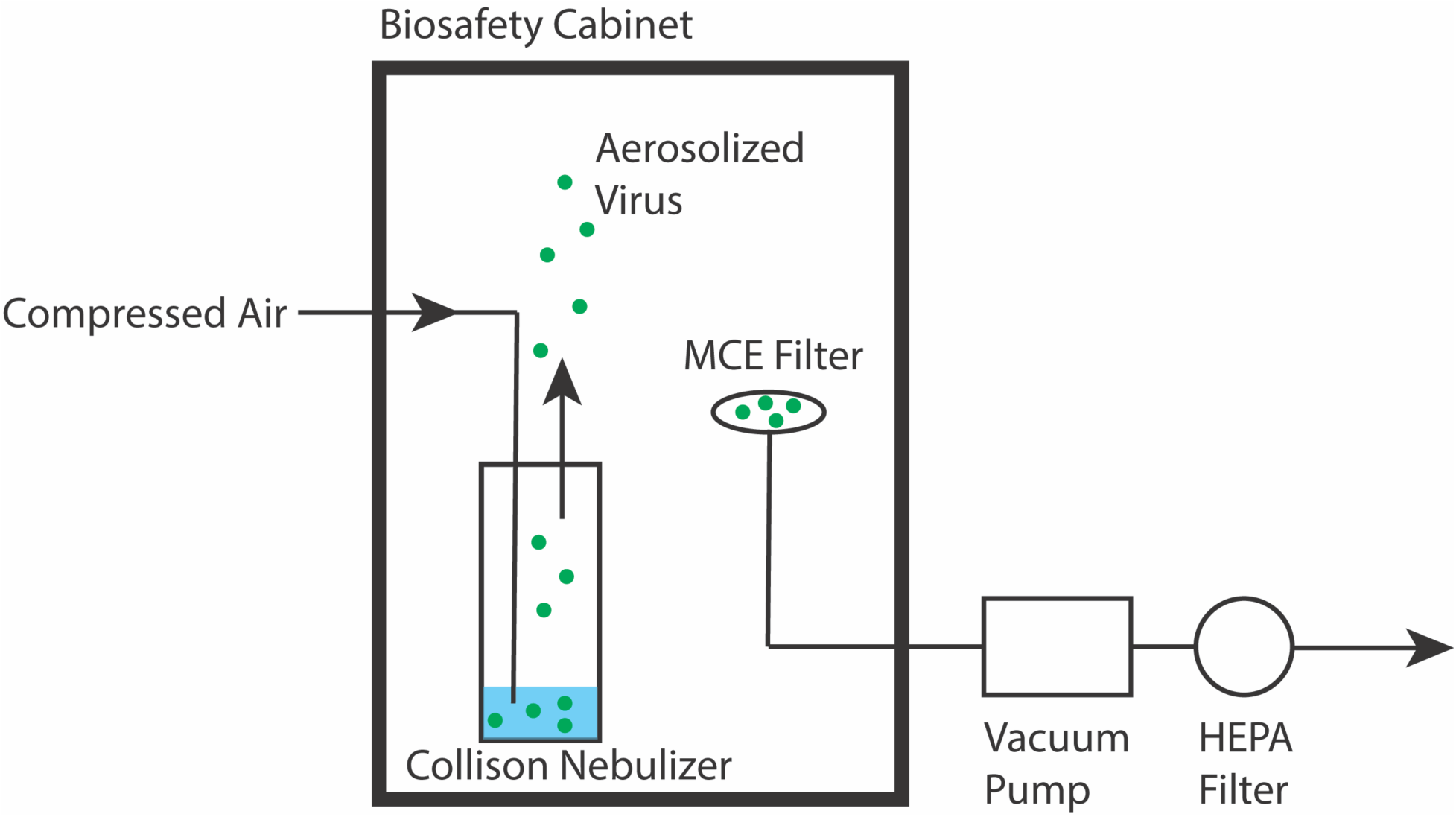
Experimental setup for aerosolized VLP.

**Figure 7:**
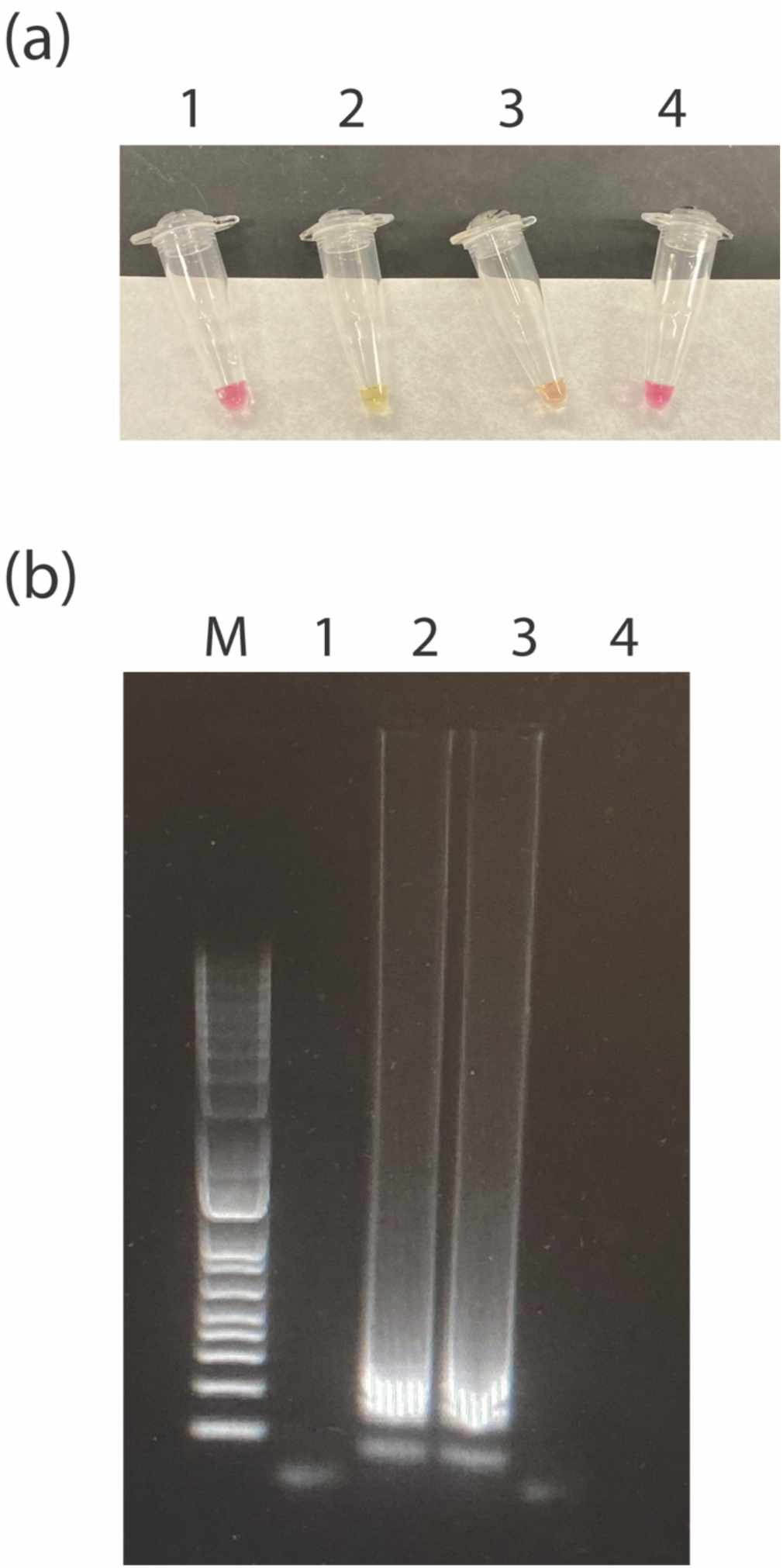
Detection of aerosolized VLP on a MCE filter. (a) Colorimetric RT-LAMP assay. (b) SYBR Green stained agarose gel of LAMP reaction products. M=Marker, 1=NTC, 2=Filter Extract, 3=Filter Extract 1/10 dilution, 4=Filter Extract 1/100 dilution.

## Conclusions

In summary, we have demonstrated the potential of using VLP for the study of airborne viral pathogens. As noted in the introduction, VLP present a number of attractive properties including their biosafety profile, facile generation, close physical resemblance to the viral pathogen of interest (SARS-CoV-2 in the present case), and readout assays by nucleic acid amplification techniques, as well as antigen-based techniques (p24 or SARS-CoV-2 spike in the present case). Moreover, different packaging vectors may be used to generate VLP to better mimic the virus of interest (e.g. that of influenza). Alternatively, the sequences for RNA genes understudy may be easily cloned into the packaging vector, thereby allowing for detection by various nucleic acid amplification protocols (e.g. in the development of devices to detect airborne viral pathogens). We suggest that this system is ideally suited for detailed studies to characterize the effects of humidity, delivery (e.g. respiration versus coughing models), suspension media (simple buffers versus artificial saliva), and membrane composition (e.g. SARS-CoV-2 spike wild type or variant or the envelope protein from another virus such as influenza hemagglutinin). We note that our groups have recently exploited the VLP model system for the development of a real-time detector of airborne viruses [28]. Taken together, we believe that the VLP-based model system will lead to new insights into the aerosolization and infection routes of numerous airborne pathogens, as well as aid in the development of devices to detect airborne viruses.

## Acknowledgements

The project described was partially supported by the National Center for Advancing Translational Sciences (NCATS), National Institutes of Health, through Grant Award Number UL1TR002003 and the UIC Chancellor’s Tranlational Research Initiatve award. The content is solely the responsibility of the authors and does not necessarily represent the official views of the NIH.

## Author Contributions

M.C., L.R. and I.P. designed the study. N.J., V.C. and V.A. contributed to the data acquisition, statistical analyses and interpretation. M.C. drafted the manuscript. I.P. and L.R. revised the article. All authors contributed to the article and approved the submitted version.

